# Endometrial zinc transporter *Slc39a10/Zip10* is indispensable for progesterone responsiveness and successful pregnancy in mice

**DOI:** 10.1101/2023.09.26.559476

**Authors:** Yui Kawata, Jumpei Terakawa, Ayuu Takeshita, Takafumi Namiki, Atsuko Kageyama, Michiko Noguchi, Hironobu Murakami, Toshiyuki Fukada, Junya Ito, Naomi Kashiwazaki

**Author notes:** Correspondence: Jumpei Terakawa, Ph.D. Laboratory of Toxicology, School of Veterinary Medicine, Azabu University, 1-17-71 Fuchinobe, Chuo-ku, Sagamihara, Kanagawa 252-5201 JAPAN, and Junya Ito, Ph.D. Laboratory of Animal Reproduction, School of Veterinary Medicine, Azabu University, 1-17-71 Fuchinobe, Chuo-ku, Sagamihara, Kanagawa 252-5201 JAPAN.

## Abstract

Zinc is a critical trace element that is important for various biological functions including male and female reproductive systems, but the molecular mechanisms that underlie fertility have been unclear. We show here for the first time that zinc signaling in the endometrial tissue is indispensable for successful embryo implantation in mice. We observed that a uterine-specific genetic deletion of *Slc39a10/Zip10*, which encodes one of the zinc transporters to elevate the cytoplasmic level of zinc, results in severe female infertility due to failure of embryo invasion into the endometrium. *Zip10* mRNA is expressed in uterine tissues, especially in the decidualizing stromal cells during embryo implantation. Absence of *Zip10* results in the suppression of zinc ion influx in the uterine stromal cells and an attenuation of progesterone – progesterone receptor signals between the epithelium and the stroma, leading to failure of embryo invasion due to sustained epithelial integrity and subsequent embryonic loss. Our findings (*i*) highlight a biological relevance of ZIP10-mediated zinc homeostatic regulation in the establishment of a successful pregnancy and (*ii*) will help to prevent infertility in humans.

## Introduction

Infertility, defined as the failure to achieve pregnancy after 12 months of regular unprotected sexual intercourse (*1*), is a global problem affecting an estimated 186 million of people of reproductive age worldwide (*2*). The causes of infertility include male factor infertility (30%), female factor infertility (35%) and combined factor infertility (20%); 15% of infertility cases remain unexplained (*3*). Growing evidence suggests that parental nutritional status affects both fertility and the offspring’s development and health (*3*, *4*). Nutrients such as folic acid, vitamins, and minerals, are important for both male and female reproduction, but the nutrient-associated mechanisms that underlie fertility have not been determined (*5*).

Zinc is an essential trace mineral that is involved in the regulation of gene expressions and protein synthesis by affecting > 300 enzyme activities (*6*, *7*). Adult humans have approx. 2–3 g of zinc in their bodies and require a zinc intake of 8 and 11 mg/day for females and males, respectively (*8*). Zinc deficiency and elevated intracellular zinc levels have a great effect on several biological processes involving the nervous, immune, and reproductive systems (*7*, *9*, *10*). Zinc deficiency during pregnancy in humans has a significant impact on maternal health and fetal and neonatal development (*11–13*). Zinc deficiency in males has been reported to cause impaired gonadal development and dysfunction (*14*). In mice, experimental zinc deficiency causes infertility, delayed fetal growth, fetal death and malformations, and delayed sexual maturation (*15–17*).

Zinc has only one valence state as the zinc ion, Zn (II), which interacts with many molecules in biological processes (*18–20*). Zinc signaling is the action of zinc ions as intracellular and extracellular signaling factors mediated by zinc-related proteins, such as zinc-finger proteins. The human body’s intracellular zinc level is tightly regulated by zinc transporters, which include two of the solute carrier protein families: SLC30A (zinc transporter, ZnT) and SLC39A (Zrt-, Irt-related protein, ZIP) (*21*). ZnT transports zinc ions from the cytoplasm to the extracellular and intracellular organelles, whereas ZIP transports zinc ions from the extracellular and intracellular organelles to the cytoplasm (*22*, *23*). In mammals, nine types of ZnT (ZnT1 to ZnT9) and 14 types of ZIP (ZIP1 to ZIP14) have been identified (*21*, *24*). These zinc transporters are expressed in a tissue- and cell-specific manner and have different physiological roles (*21*, *25*, *26*). As zinc homeostasis has a great impact on reproductive outcomes, zinc transporters are suspected to play one or more important roles in the establishment and maintenance of pregnancy through the regulation of zinc dynamics, but the details remain elusive.

In this study, we focused on SLC39A10/ZIP10, which localizes to the cell membrane and is responsible for the influx of zinc ions into the cytoplasm (*27*). It has been reported that ZIP10 plays an important role in regulating homeostasis and functions of B lymphocytes (*28*, *29*), suggesting that ZIP10 is necessary for B cell-mediated acquired immunity. As demonstrated by the data in the gene annotation portal bioGPS (http://biogps.org/) (*30*), *ZIP10* expression is relatively high in the human and mouse uterus and placenta. To clarify the role of ZIP10 during pregnancy, we have generated uterine-specific *Zip10*-deficient mice, and we observed that these females are extremely infertile. Our findings help clarify the precise mechanism of infertility caused by lack of *Zip10* in the endometrium.

## Materials and methods

All chemicals and reagents were purchased from Sigma-Aldrich (St. Louis, MO) unless otherwise stated. All animal procedures were approved by the Ethical Committee for Vertebrate Experiments at Azabu University (ID#200312-24). All experiments were conducted in accord with the relevant guidelines and regulations, including the Animal Research: Reporting of In Vivo Experiments (ARRIVE) guidelines.

### Animals

The mice were housed in the barrier facility at Azabu University. The following three mouse strains were used: *Pgr^Cre/^*^+^ mouse <B6.129S(Cg)-*Pgr^tm1.1(cre)Shah^*/AndJ, JAX: 017915> (*31*), *Slc39a10/Zip10^flox/flox^* (*Zip10^f/f^*) mouse <*Slc39a10^tm1.1Tfk^*> (*32*), and *Ltf^iCre/+^* mouse <*Ltf^tm1(icre)Tdku^/J,* JAX: 026030> (*33*). All strains were maintained on the C57BL/6J background purchased from Jackson Laboratory Japan (Kanagawa, Japan). All mice were fed *ad libitum* under a 12-h light/12-h dark photocycle at 23 ± 2°C.

Uterine-specific *Slc39a10*/*Zip10*-deficient (*Zip10* conditional knockout [cKO]) mice were generated by several crossings of the *Pgr^Cre/+^* strain with the *Zip10^flox/flox^* strain (Suppl. Fig. S1). The genotype of the mice was confirmed by the polymerase chain reaction (PCR) of genomic DNA obtained from tail tissue (Suppl. Fig. S1A).

The PCR primers used for genotyping were 5′-AGTTATTGCTGCCCAGTTGC-3′, 5′-CCCTTTCTCATGGAGATCTGTC-3′, and 5′-GCGCTAAGGATGACTCTGGTC-3′ for *Pgr^Cre^*; and 5′-CAAGGCCAGCCAAAATTCTA-3′, 5′-GCTTTCCTCCCATCCTGATT-3′, and 5′-GTGGCATGCGTGGAAGTTAG-3′ for *Zip10^flox^*. Uterine epithelial-specific *Slc39a10*/*Zip10*-deficient (*Zip10* ecKO) mice were generated by crossing the *Ltf^iCre/+^* strain with the *Zip10^flox/flox^* strain as described (*34*). Sexually mature (> 8 weeks old) *Pgr^Cre/+^*; *Zip10^flox/flox^*and *Pgr^Cre/+^*; *Zip10^flox/−^* female mice were used as *Zip10* cKO mice. *Pgr^+/+^*; *Zip10^flox/flox^*and *Pgr^+/+^*; *Zip10^flox/−^* females were used as the control mice, since unexpected germline recombination was sometimes identified in *Pgr^Cre/+^* lines as reported (*35*) (Suppl. Fig. S1B).

Females were mated with a fertile wildtype male. The day when the vaginal plug was first observed was designated as the first day of pregnancy (D1). Mice were euthanized by cervical dislocation after a combination anesthetic with 0.75 mg/kg of medetomidine, 4.0 mg/kg of midazolam, and 5.0 mg/kg of butorphanol at indicated day(s) of pregnancy (*36*). Uterine and ovarian tissues harvested at indicated day(s) of pregnancy were fixed with 4% paraformaldehyde (PFA) solution in phosphate-buffered saline (PBS) for histological study, fixed with G-Fix (Genostaff, Tokyo) for i*n situ* hybridization (ISH) and snap-frozen and kept at −80 °C until used for RNA extraction.

The implantation sites were visualized by an intravenous injection of 0.1 mL of 1% Chicago blue dye dissolved in saline as described (*37*). After female mice were mated with a vasectomized male, artificial decidualization was performed by the administration of intraluminal injections of 0.02 ml of sesame oil as described (*38*).

In order to control the embryo implantation process by an administration of exogenous hormone, the ovaries of control and *Zip10* cKO females were removed at D4, followed by the insertion of a Silactic implant (20 mm long × 2.0 mm inner dia., 3.0 mm outer dia., Silascon 100-2N, Kaneka Medix, Osaka, Japan) containing progesterone (P4) powder under the dorsal skin (Suppl. Fig. S3A). Two days later, a single subcutaneous injection of 17β-estradiol (E2, 25 ng/head) was performed to induce embryo implantation. Uterine tissues were collected 3 days after the E2 injection to verify successful embryo implantation and decidualization.

### Measurement of serum hormone levels

Blood samples from indicated mice were collected through cardiac puncture under combination anesthetic on D5, 6, 8 and 10 of pregnancy. Serum was separated by centrifugation (4°C, 7,000 g, 7 min) and stored at −80°C until analysis. The serum concentrations of P4 and E2 were measured by enzyme-linked immunosorbent assays as described (*39*).

### Histology

Fixed uterine and ovarian tissues were paraffin-embedded, and the paraffin sections (6 µm) were stained with hematoxylin and eosin (H&E).

### In situ hybridization (ISH)

In situ hybridization (ISH) was performed as described with some modifications (*32*). Briefly, fixed uterine tissues were paraffin-embedded, and the paraffin sections (6 µm) were mounted on MAS-coated slides (Matsunami Glass Industries, Osaka, Japan) under RNase-free conditions. Sense or antisense digoxigenin (DIG)-labeled RNA probes for *Slc39a10*/*Zip10* were purchased from Genostaff (Tokyo). The sections were deparaffinized, rehydrated, and post-fixed in 10% neutral buffered formalin (NBF) for 30 min at 37°C, followed by treatment with 0.2% hydrogen chloride and 5 µg/mL proteinase K (FujiFilm Wako Pure Chemicals, Osaka, Japan) for 10 min at 37°C, respectively. Hybridization was performed with DIG-labeled probes (300 ng/300 µL/ slide) in a humidified chamber at 60°C overnight.

The slides were washed after hybridization, then treated with blocking reagent (Genostaff) for 15 min and with alkaline phosphatase-conjugated anti-DIG antibody (1:2,000; Roche Diagnostics, Basel, Switzerland) for 1 h at room temperature. The signals were detected by 4-nitro-blue tetrazolium/5-bromo-4-chloro-3-indolyl phosphate (NBT/BCIP, Roche Diagnostics) in a humidified container for 72 h at 4°C. The sections were counterstained with Kernechtrot solution (Muto Pure Chemicals, Tokyo). Signals detected by the sense probe were used as a control for background levels.

### Immunohistochemistry (IHC) and immunofluorescence

Fixed tissues were paraffin-embedded, and the paraffin sections (6 µm) were deparaffinized, hydrated, and used for antigen retrieval by autoclaving in 10 mM sodium citrate buffer (pH = 6.0) for 5 min. For immunohistochemistry (IHC), the sections were further incubated in 3% hydrogen peroxide diluted with methanol for 15 min. After blocking with a non-specific staining blocking reagent (X0909, Dako, Carpinteria, CA), the slides were incubated with primary antibodies (shown in Suppl. Table S1) overnight at 4°C. The same slides were subjected to incubation with the Histofine mouse stain kit (Nichirei Biosciences, Tokyo) for 1 h. Signals were visualized by 3,3’-diaminobenzidine tetrahydrochloride (DAB) and counterstained with hematoxylin.

For the immunofluorescence evaluation, the slides were incubated with Alexa Fluor 488-conjugated secondary antibodies (Jackson Immuno Research Laboratories, West Grove, PA) for 1 h, and mounted with ProLong Glass Antifade Mountant with NucBlue Stain (P36981, Thermo Fisher Scientific, Waltham, MA). Micrographs were captured by PROVIS AX80 microscopy (Olympus, Tokyo) or BZ-X700 microscopy (Keyence, Osaka, Japan). All signals were detected under the same lighting conditions for the control and *Zip10* cKO groups.

### RNA extraction and quantitative PCR (qPCR)

Total RNA was isolated using the RNeasy Plus Mini kit (Qiagen, Venlo, Netherlands) according to the manufacturer’s instructions. After the total RNA concentration was measured using a Nanodrop system (ND-1000, Thermo Fisher Scientific), 1 µg of total RNA in each sample was reverse-transcribed with SuperScript III Reverse Transcriptase (#18080-044, Thermo Fisher Scientific) using oligo-dT primer (#3805, Takara, Shiga, Japan). A quantitative (q)PCR was performed on a Thermal Cycler Dice Real Time System (Takara) with the use of Power SYBR Green Master Mix (A25742, Thermo Fisher Scientific). The primer sequences in this study are shown in Supplemental Table S2. The relative expression values of the target transcripts were evaluated using the ΔΔCt relative quantification method (*40*) with normalization to *Gapdh*. The average values of control samples were set as 1.0.

### Isolation and culture of primary uterine stromal cells

For the isolation and culture of primary uterine stromal cells, uterine stromal cell isolation and culture were performed as described with minor modifications (*41*). Briefly, uterine horns collected on day 4 of pregnancy were rinsed with phenol red-free Hank’s balanced salt solution (HBSS)(−) (#085-09355, FujiFilm Wako Pure Chemicals) containing 1× antibiotic-antimycotic solution (#161-23181, FujiFilm Wako Pure Chemicals) for the removal of blood and trimmed fat and mesometrial tissues. Uterine tissues were cut into 3- to 4-mm lengths with sterile scissors and incubated in phenol red-free HBSS(−) containing dispase (#17105-041, Thermo Fisher Scientific) and pancreatin (#P3292) at 4°C for 1 h, followed by incubation at room temperature (RT) for 1 h and at 37.5°C for 10 min. The enzyme reaction was stopped by the addition of fetal bovine serum (FBS) (Biowest, Nuaillé, France).

Uterine pieces were washed with phenol red-free free HBSS(−) to remove epithelial cells and further incubated in phenol red-free free HBSS(−) with collagenase type III (#CLS3, Worthington, Lakewood, NJ) at 37.5°C for 30 min. After the addition of FBS, the uterine pieces were washed with phenol red-free free HBSS(−) and passed through a 70-μm cell strainer (Corning, Glendale, AZ). Isolated uterine stromal cells were concentrated by centrifugation (900 g, 7 min), inoculated into 35-mm glass-bottomed dishes (Matsunami Glass Industries), and incubated with D-MEM/Ham’s F-12 culture medium (#045-30665, FujiFilm Wako Pure Chemicals) containing 10% charcoal-stripped FBS (CS-FBS) (Cytiva, Tokyo) and 1× penicillin/streptomycin solution (#168-23191, FujiFilm Wako Pure Chemicals) at 37.5°C under 5% CO2. The culture medium was changed 1 h after inoculation, and adherent cells were continuously incubated for 24 h.

For the induction of *in vitro* decidualization, the medium was replaced with D-MEM/Ham’s F-12 containing 2% CS-FBS, 1 nM E2, and 1 μM P4. The detection of intracellular zinc levels by FluoZin-3AM (Thermo Fisher Scientific) and fluorescent immunocytochemistry (FICC) for cultured stromal cells was performed 48 h after the medium replacement. Alkaline phosphatase (ALP) staining with BCIP/NBT Color Development Substrate solution (#S3771, Promega, Madison, WI) was performed 96 h after the medium replacement as described (*42*, *43*).

### Statistical analysis

All of the data are presented as the mean ± standard error of mean (SEM). Differences in the data were examined with the use of Student’s *t*-test or the Mann‒Whitney U-test. A *p*-value <0.05 was considered significant.

## Results

### Zip10 expression in the pregnant mouse uterus

We first examined the *Zip10* expression in the pregnant mouse uterus. *Zip10* mRNA expression was detected in the uterine epithelium at the first day (D1) of pregnancy (Fig. 1). A relatively strong expression of *Zip10* in stromal cells was observed on D5, when the embryos undergo an attachment reaction to the luminal epithelium. Stromal expression of *Zip10* was observed in decidualizing stromal cells, and secondary decidualized cells were clearly positive for *Zip10* on D8 as shown in Figure 1.

**Fig. 1.**
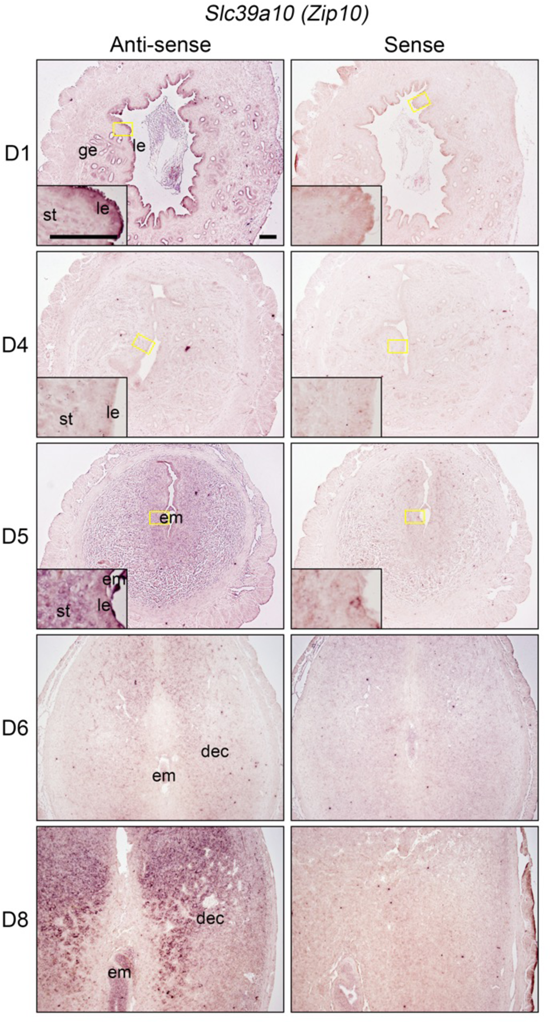
Spatiotemporal expression of *Slc39a10/Zip10* mRNA in the pregnant mouse uterus. Signals (*dark purple*) were detected by an anti-sense probe (*left*). The sense probe (*right*) shows the background signal. *Slc39a10/Zip10* mRNA was detectable in the luminal and glandular epithelium at day 1 of pregnancy (D1), in differentiating stromal cells at D5, and in decidual cells at D8. Scale bar: 100 µm. dec: decidua, em: embryo, ge: glandular epithelium, le: luminal epithelium, st: stroma.

### Pregnancy failure of the uterine-specific Zip10-deficient females

To determine whether the deletion of the *Zip10* gene in the uterine tissue affected the pregnancy outcomes in the mice, we generated uterine-specific *Zip10*-deficient mice (*Zip10* cKO) driven by *Pgr^Cre^*, which targets directed recombination in the female reproductive tracts (*31*, *35*) (Suppl. Fig. S1). We crossed the control and *Zip10* cKO females with sexually mature wildtype males. Control females (n = 11) gave birth to an average of 6.8 ± 0.5 pups, whereas only five out of the 23 *Zip10* cKO females gave birth; they produced a total of 13 pups (Fig. 2A). The average number of pups produced by the *Zip10* cKO females was 0.6 ± 0.3 (Fig. 2A), demonstrating that the *Zip10* cKO mice exhibited severe infertility. We then examined the pregnant uteri on D5, D6, D8, and D10 to determine when pregnancy failure occurred (Fig. 2B).

**Fig. 2.**
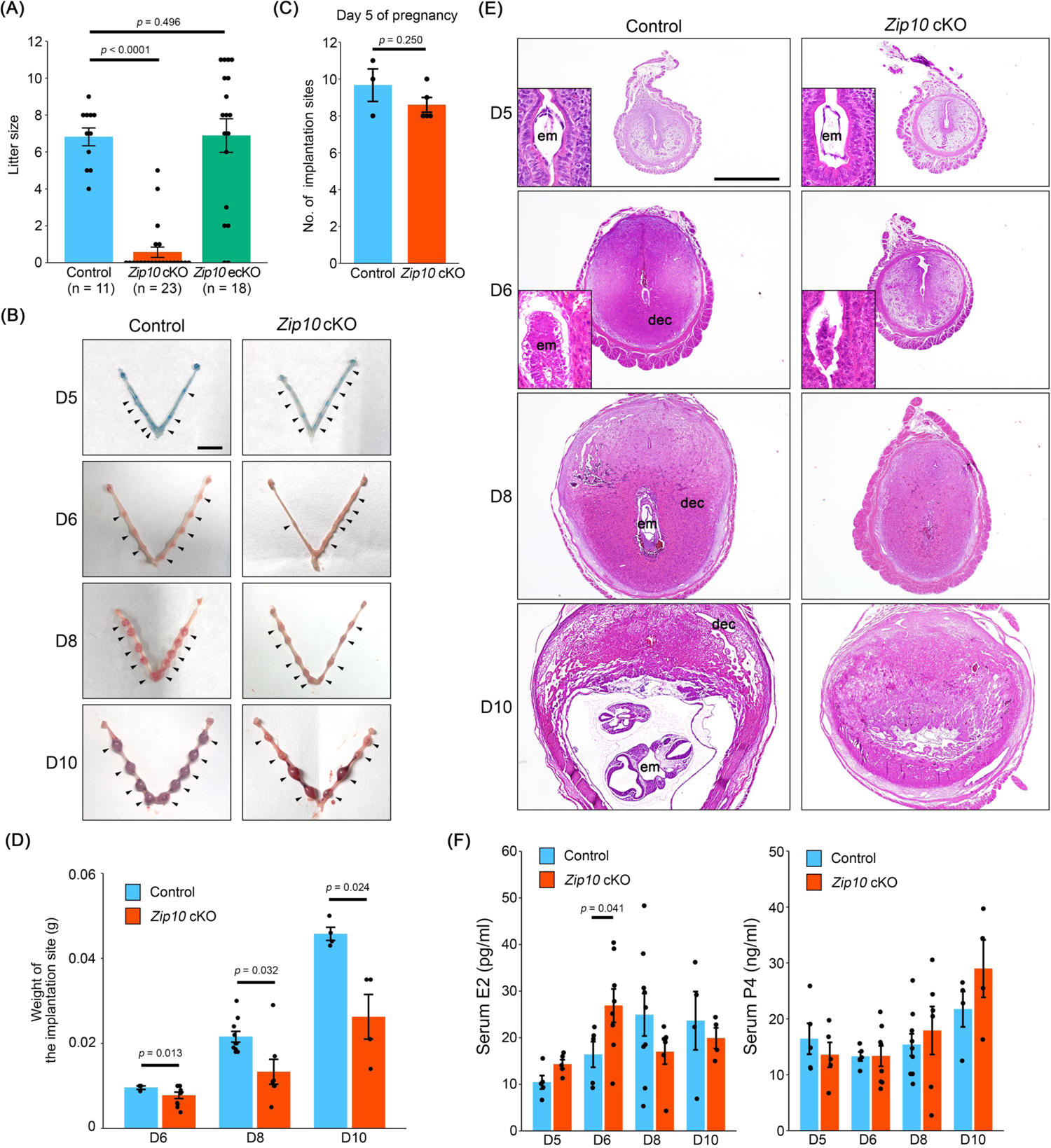
Failure of pregnancy maintenance in uterine-specific *Slc39a10/Zip10*-deficient females. **(a)** Pregnancy outcomes from control, *Zip10* cKO, and *Zip10* ecKO females. **(b)** Representative uteri from control and *Zip10* cKO females from D5 to D10. Implantation sites are shown by *arrowheads* (visualized by a blue dye injection) on D5. Scale bar: 1 cm. **(c)** The number of implantation sites were comparable between the control and *Zip10* cKO females on D5. **(d)** The average wet weight of each implantation site was significantly reduced in the *Zip10* cKO mice on D6, D8, and D10. **(e)** Representative images of uterine cross-sections stained by H&E. Scale bar: 1 mm. le: luminal epithelium; dec: decidua. **(f)** Serum levels of 17β-estradiol (E2) and progesterone (P4) from D5 to D10.

On D5, the blue-dye injections visualized the implantation sites in *Zip10* cKO mice as well as the control mice, suggesting the number of implantation sites was comparable between the control and Z*ip10* cKO groups (Fig. 2B,C). From D6 to D10 however, the wet weight of each implantation site became significantly lower in the Z*ip10* cKO females compared to the control females (Fig. 2D). Histological observations revealed that on D5, (*i*) embryos were positioned at the anti-mesometrial side of the slit-like uterine lumen, and (*ii*) no apparent difference in the positions between the Z*ip10* cKO and control groups (Fig. 2E). Nevertheless, on D6 the uterine lumen was not closed and decidualization was not sufficiently induced in the Z*ip10* cKO females, with the result that embryos did not invade the endometrium (Fig. 2E). In the Z*ip10* cKO females, embryonic loss was observed as early as D8, and most of the embryos were regressed and resorbed by D10 (Fig. 2B,E).

Since both 17β-estradiol (E2) and progesterone (P4) are required to induce optimal decidualization, we measured the serum E2 and P4 levels in control and Z*ip10* cKO females. On D6 of pregnancy, the E2 levels were higher in the Z*ip10* cKO females compared to the control females, whereas the P4 levels were comparable between the control and Z*ip10* cKO females (Fig. 2F). No significant difference was observed in ovarian morphology; both groups of females developed multiple corpus lutea (Suppl. Fig. S2). To test the possibility that higher E2 levels affect the pregnancy outcome in Z*ip10* cKO females, we performed ovariectomies and exogenous E2 and P4 administrations to control the hormone levels and timing of embryo implantation (Suppl. Fig. S3A). The experimental procedure in this study induced a one-day delay in the embryo implantation and decidualization process. The control females exhibited the successful embryo implantation and decidualization, but the Z*ip10* cKO females failed to undergo optimal decidualization despite the control of the hormone levels (Suppl. Fig. S3B,C). These results suggest that the embryonic loss in Z*ip10* cKO females is due mainly to the functional defect of the endometrium, which manifests itself by D6 of pregnancy.

Since *Zip10* cKO females derived from the *Pgr^Cre^*strain lose ZIP10 expression in the whole endometrium, we established uterine epithelial-specific *Zip10*-deficient (*Zip10* ecKO) mice by using the *Ltf^iCre/+^* strain. The *Zip10* ecKO females were fertile; the average litter size was 6.9 ± 0.9 (n=18) (Fig. 2A) and there was no evidence of embryonic loss, indicating that stromal ZIP10, but not epithelial ZIP10, is indispensable for successful pregnancy.

### Abnormal progesterone receptor expression in the Zip10 cKO uterine epithelium

To further investigate the pregnancy failure in *Zip10* cKO females, we performed an immunofluorescence evaluation for the epithelial marker CDH1 (also known as E-cadherin), in the uterine tissue during embryo implantation (Fig. 3A). In the controls, luminal epithelial cells were removed from D5 to D6, and embryos were placed directly in contact with decidualized stromal cells (Fig. 3A). In contrast, we observed on D6 that in the *Zip10* cKO females the luminal epithelial cells had remained intact (Fig. 3A). Since the cessation of luminal epithelial cell proliferation is essential for embryo implantation and since aberrant epithelial proliferation blocks the embryo implantation process (*44*), we examined the cell proliferation status by performing MKI67 staining on D5 (Fig. 3B). The cell proliferation status of the luminal epithelium was comparable between the control and *Zip10* cKO groups. On the other hand, the proliferation of subluminal stromal cells was increased in the *Zip10* cKO compared to the control groups (Suppl. Fig. S4)

**Fig. 3.**
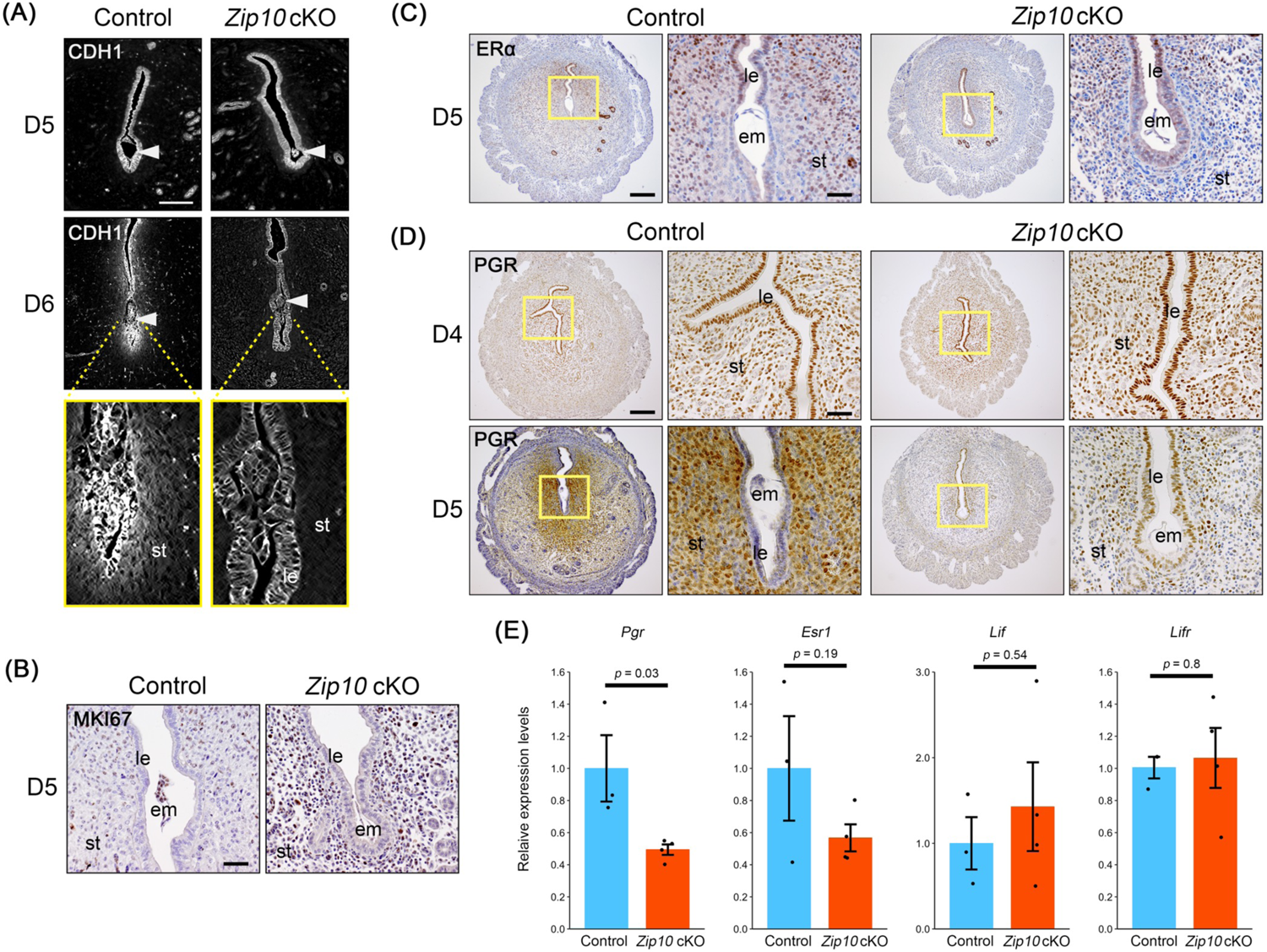
Extraction of luminal epithelial cells surrounding the embryo was not induced in *Zip10* cKO mice. **(a)** Immunofluorescence of CDH1 in sections of implantation sites on D5 and D6. The embryo (*arrowhead*) directly contacts subepithelial stroma in the controls on D6, but not in the *Zip10* cKO mice. Scale bar: 200 µm. le: luminal epithelium; st: stroma. (b–d) Immunostaining for MKI67 **(b)**, ERα **(c)**, and PGR **(d)**. Epithelial PGR expression is absent in the controls on D5 but was sustained in the *Zip10* cKO mice. Scale bar: 50 µm (b, right panels of c and d) and 200 µm (c and d: left panels). le: luminal epithelium; st: stroma; em: embryo. **(e)** Relative mRNA expressions of *Pgr, Esr1*, and the implantation-related factors *Lif* and *Lifr*.

We further examined the expression of estrogen receptor alpha (ERα, ESR1) and progesterone receptor (PGR) by IHC during early pregnancy (Fig. 3C,D). No remarkable difference in ERα expression was observed on D5 (Fig. 3C). PGR expression was switched from the luminal epithelium to the stroma in the controls from D4 to D5, but PGR was continuously expressed in the luminal epithelium in the *Zip10* cKO group on D5 (Fig. 3D). The results of the qPCR analysis confirmed that the overall expression of *Pgr* mRNA was decreased in the *Zip10* cKO group on D5, which is consistent with the weak expression of PGR in stromal cells (Fig. 3D,E). Not only *Esr1* mRNA but also the expressions of leukemia inhibitory factor (*Lif*), a key regulator of embryo attachment (*34*) and its receptor, *Lifr,* were not significantly different between the control and *Zip10* cKO groups on D5 (Fig. 3E). These results suggest that progesterone receptor signaling appears to be disrupted in *Zip10* cKO mice, leading to the prevention of on-time embryo implantation.

### Defects of progesterone receptor signaling in the Zip10 cKO uterus

Since PGR expression was abnormally sustained in the luminal epithelium in the *Zip10* cKO group during the embryo implantation process, we investigated the expression of downstream targets of PGR. The results demonstrated that preceded by small implantation sites in *Zip10* cKO at D6, the expression of well-known genes related to the decidualization, including *Hoxa10*, *Bmp2*, and *Wnt4* (*41*, *45*) were significantly decreased in the *Zip10* cKO uteri compared to the control uteri on D5 (Fig. 4A). Indian hedgehog (*Ihh*), which is expressed in the uterine epithelium under the control of PGR (*46*), was significantly upregulated in the *Zip10* cKO group compared to the controls (Fig. 4A). Activated IHH acts on patched receptor 1 (*Ptch1*) in neighboring cells to remove smoothened (*Smo*) inhibition, activating the glioma-associated oncogene homolog 1 (*Gli1*) and inducing factors such as chicken ovalbumin upstream promoter transcription factor II (COUP-TFII, encoded by *Nr2f2*) (*47*).

**Fig. 4.**
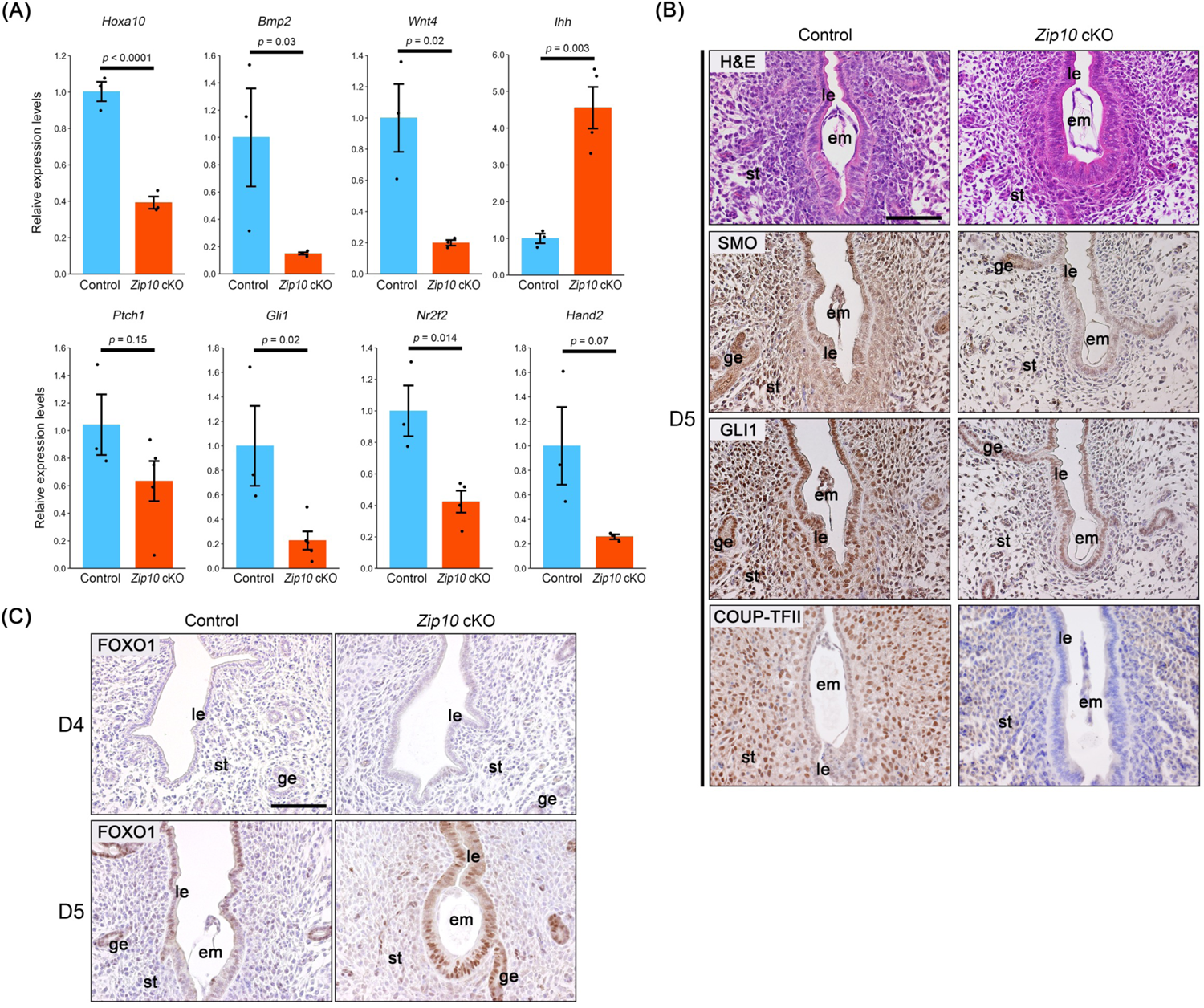
P4-PGR target genes are disrupted in *Zip10* cKO mice. **(a)** Relative mRNA expressions of decidual marker genes and P4-PGR target genes. **(b)** H&E staining and immunostaining for SMO, GLI1, and COUP-TFII. These expressions were reduced in the *Zip10* cKO uteri compared to the controls. Scale bar: 100 µm. **(c)** Immunostaining for FOXO1. FOXO1 was expressed in the uterine epithelium at the timing of embryo implantation in the Zip10 cKO mice. Scale bar: 100 µm. em: embryo, ge: glandular epithelium, le: luminal epithelium, st: stroma.

Intriguingly, the expressions of *Gli1* and *Nr2f2* mRNA were significantly decreased on D5 in *Zip10* cKO group compared to the controls (Fig. 4A). These results were further confirmed by the immunostaining; the expressions of SMO, GLI1, and COUP-TFII were markedly decreased in the pregnant uteri of *Zip10* cKO mice, especially in the stroma (Fig. 4B, Suppl. Fig. S5). The expression of a stromal PGR mandated factor, i.e., heart and neural crest derivatives expressed 2 (*Hand2*) (*44*) was also downregulated in the *Zip10* cKO uteri (Fig. 4A). Taken together, these results indicated that progesterone receptor signaling was disrupted in the *Zip10* cKO mice.

It has been demonstrated that epithelial PGR stability is controlled by the transcription factor, forkhead box O1 (FOXO1), based on the observation that FOXO1 ablation resulted in an increase in PGR signaling, as PGR expression was retained in the uterine epithelium during the window of receptivity (*48*). We therefore analyzed the expression of FOXO1 and found that it was expressed in the uterine epithelium of both the control and *Zip10* cKO mice (Fig. 4C), suggesting that FOXO1 was not a primary target of ZIP10 deficiency.

We also detected the expression of *Epas1* (also known as hypoxia-inducible factor 2 alpha [*Hif2α*]), which is expressed in the subepithelial stroma and regulates embryo invasion (*49*, *50*). The expression of *Epas1* was comparable between the control and *Zip10* cKO uteri on D6 (Suppl. Fig. S6).

### Zinc depletion and decidualization in Zip10 cKO uterine stromal cells

We next investigated whether decidualization is inducible in the *Zip10* cKO uterine stroma. Interestingly, decidualization was partially induced by the intrauterine infusion of sesame oil in *Zip10* cKO mice (Fig. 5A). We thus isolated stromal cells from both control and *Zip10* cKO mice on D4 and performed *in vitro* decidualization using E2 and P4 (*41*) (Fig. 5B).

**Fig. 5.**
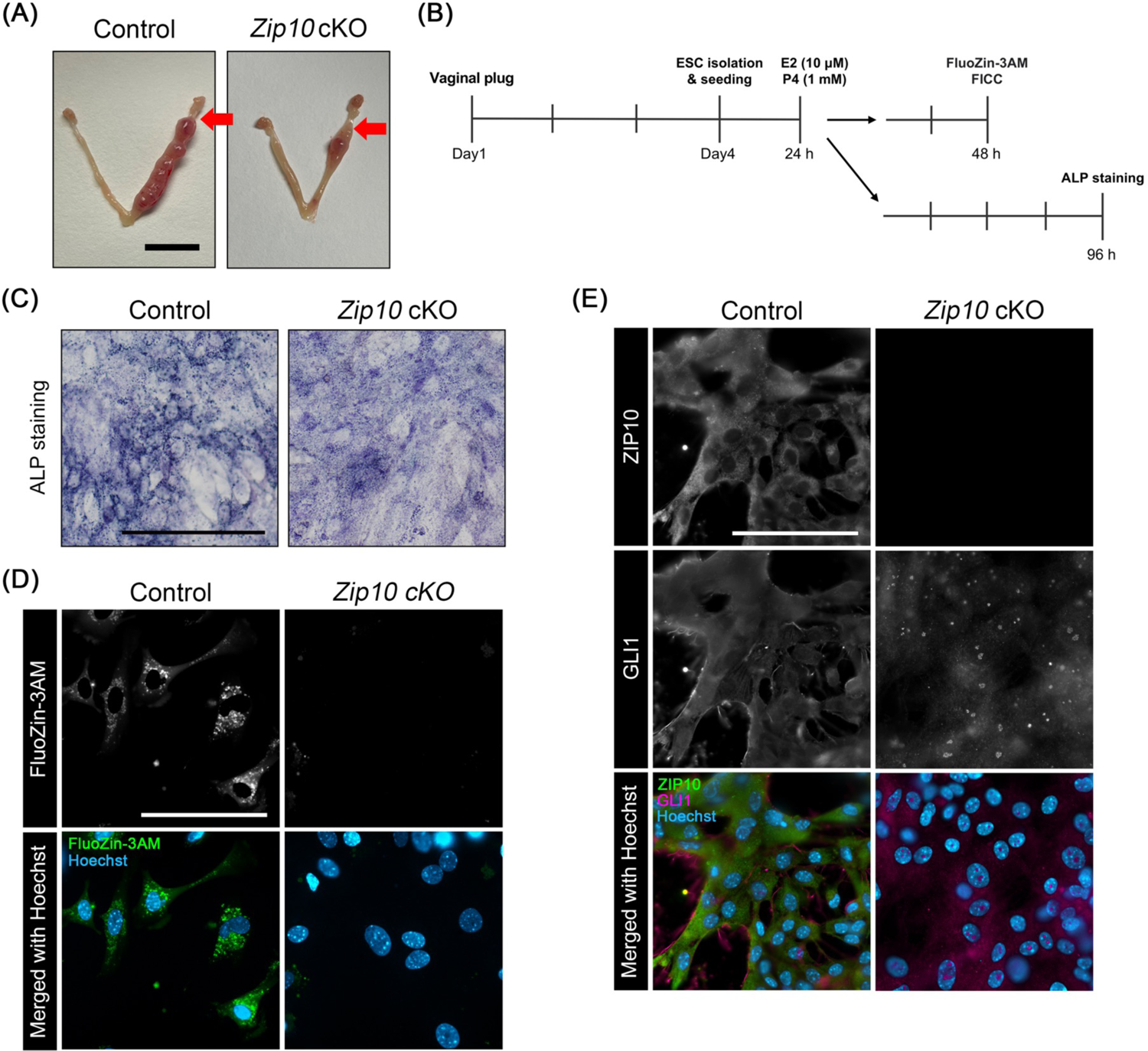
Attenuation of *in vivo* decidual reaction in *Zip10* cKO mice. **(a)** Representative images showing the gross morphology of artificial decidualization from control and *Zip10* cKO mice. *Red arrows:* the site of sesame oil injection. Scale bar: 1 cm. **(b)** Experimental procedure for endometrial stromal cell (ESC) isolation and *in vitro* decidualization. **(c)** Alkaline phosphatase (ALP) staining for decidualized cultured stromal cells. Scale bar: 100 µm. **(d)** Staining with FluoZin-3AM, a zinc ion indicator, in cultured stromal cells. No positive signal for FluoZin-3AM was observed in the cytoplasm of *Zip10* cKO stromal cells. Scale bar: 100 µm. **(e)** Fluorescent immunocytochemistry (FICC) for ZIP10 and GLI1. GLI1 was detected in the cytoplasm of control stromal cells but was localized in the nucleus in *Zip10* cKO stromal cells. Scale bar: 100 µm.

The results demonstrated that decidualization can be induced by isolated primary stromal cells from *Zip10* cKO mice as well as control mice by assessing positive ALP staining (Fig. 5C), suggesting that stromal cells have the ability to differentiate.

To address whether the deletion of *Zip10* led to intracellular zinc depletion, we examined the intracellular zinc level by using a zinc ion indicator, FluoZin-3AM. FluoZin-3AM fluorescence was detected in the stromal cells of control females, but not those of *Zip10* cKO females (Fig. 5D). We confirmed that ZIP10 was absent in *Zip10* cKO stromal cells (Fig. 5E).

We also observed the clear differentiation of GLI1 protein localization between the control and *Zip10* cKO stromal cells; GLI1 was localized in the cytoplasm in the control cells but in the nucleus of *Zip10* cKO cells, supporting the concept that GLI1 localization is regulated by the intracellular zinc level.

## Discussion

Our present findings demonstrate that intracellular zinc, regulated by the endometrial zinc transporter ZIP10, is essential for successful embryo implantation and placentation in mice. In particular, the results revealed that an intracellular zinc deficiency due to a lack of ZIP10 impairs the P4 responsiveness of the endometrium, resulting in both the failure of degradation for the luminal epithelium and severe embryonic loss (Fig. 6).

**Fig. 6.**
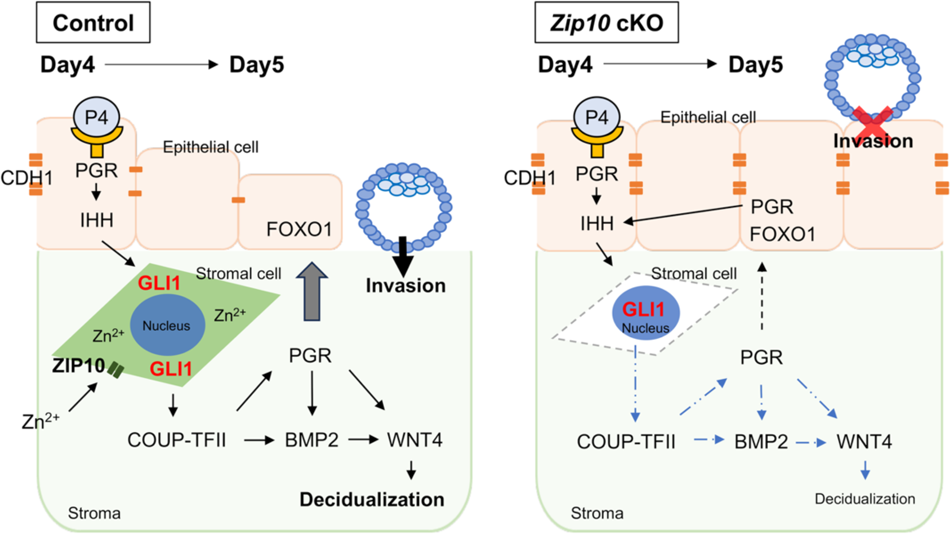
Model of embryo invasion regulated by zinc dynamics though ZIP10. Zinc ion regulates P4-PGR signaling during embryo implantation. In *Zip10* cKO mice, zinc ion is not incorporated into the uterine cells, leading to the nuclear localization of stromal GLI1 and attenuation of P4-PGR signaling. This results in incomplete decidualization and a failure of the elimination of epithelial cells that allows embryonic invasion, and thus subsequent embryonic loss.

In this study, the deletion of ZIP10 by *Pgr^Cre^* manifested a phenotype as early as D5 of pregnancy as abnormally sustained PGR expression in the luminal epithelium with a decreased expression of most P4 target genes, although the morphology of the uterine tissues of the *Zip10* cKO and control mice was comparable. The initial reaction of embryo implantation, including embryo apposition and attachment, was not disrupted in the *Zip10* cKO mice, as aligned implantation sites was clearly visualized by blue dye and the number of implantation sites was not decreased in the *Zip10* cKO mice on D5 (Fig. 2B,C,E). This is further confirmed by our observation that the expressions of *Lif* and *Lifr*, which are essential factors for embryo attachment (*51*), were not significantly different between the *Zip10* cKO and control mice (Fig. 3E). On D6, failure of the embryo implantation process — more specifically failure of embryo invasion — was elicitable, as shown by our finding that embryos were not in direct contact with stromal cells in the *Zip10* cKO mice (Fig. 3A). This failure was most likely associated with a weak decidualization reaction and subsequent embryonic loss after D6 (Fig. 2D,E).

Genetic recombination using *Pgr^Cre^* may target not only the uterus but also the ovaries and pituitary gland. In fact, it has been reported that *Lgr5* gene deletion by *Pgr^Cre^* results in severe infertility due to a disruption of P4 production in the corpus luteum (*52*). Unlike the *Pgr^Cre^* strain used in that study, the *Pgr^Cre^* strain used in our present investigation was based on ires-dependent Cre recombinase expression and was considered to have a minor genetic defect in the newly formed corpus luteum (*35*), but the potential genetic defect still elicits the phenotype. In our study, at least, no P4 depletion was observed in the *Zip10* cKO mice during early pregnancy (Fig. 2F). To exclude the possibility that a pituitary-ovarian axis defect caused the infertile phenotype in the *Zip10* cKO mice, we used a delayed implantation model. In mice, pregnancy can be maintained by the administration of external hormones after the removal of ovaries in the delayed implantation model (*53*, *54*). Our results demonstrated that the imbalance of hormones was not the main cause of pregnancy failure in the *Zip10* cKO mice. The E2 levels were 1,000 times lower than those of P4, which may have resulted in the differences in variation among the individual mice.

We also observed that the deletion of ZIP10 in the endometrium, especially stromal cells, was not directly linked to the functional loss for decidualization, because weak decidualization reaction was observed in the *Zip10* cKO uteri (Fig. 2E) and the partial deciduoma could be induced after an intrauterine oil infusion (Fig. 5A). In addition, after the induction of *in vitro* decidualization, isolated stromal cells from *Zip10* cKO mice exhibited ALP activity, which is a well-known marker of stromal differentiation (*41*) (Fig. 5B,C). More interestingly, the uterine epithelial loss of *Zip10* (*Zip10* ecKO) did not result in any reproductive failure (Fig. 1A), indicating that the cause of pregnancy failure observed in the *Zip10* cKO mice is most likely due to a failure of epithelial-stromal interaction during early pregnancy.

The pre-receptive mouse uterus (D1 to D3) is under an E2-dominant state, with proliferation of epithelial cells (*55*). P4 signal, mediated by both epithelial and stromal PGR, counteracts E2, and it is required for implantation of embryos in the receptive uterus (*56*). IHH, a secreted morphogen, is transiently expressed in the preimplantation uterine epithelium by P4-PGR (*46*, *57*), and its expression is also reported to be enhanced by LIF stimulation (*58*). Secreted IHH from the uterine epithelium binds to the canonical IHH receptor PTCH1 in the underlying stroma, resulting in the de-repression of SMO and the activation of GLI, which is a zinc finger transcription factor.

Activation of the IHH pathway leads to a promotion of the expression of COUP-TFII (also known as NR2F2), which subsequently upregulates the stromal PGR expression and induces BMP2, WNT4, and HAND2, which are essential factors for decidualization (*56*). HAND2 is also a stromal P4-PGR target transcription factor that normally inhibits the expression of several members of the fibroblast growth factor (FGF) family that promotes uterine epithelium proliferation (*44*, *56*). HOXA10 is another stromal P4-PGR target transcription factor involved in decidualization (*59*). In the present study, the expressions of these P4-PGR target genes were disrupted; most of them were downregulated, with the exception of *Ihh* (Fig. 4). Increased expression of *Ihh* is closely related to the sustained PGR expression in the luminal epithelium. Under normal conditions, PGR expression shifts from the uterine epithelium to the uterine stroma in the progression of the embryo implantation process (*60*).

Wetendorf *et al*. reported that a constitutive expression of PGR in the luminal epithelium of transgenic strains resulted in the failure of embryo attachment, and it was concluded that a loss of epithelial PGR expression is necessary for successful embryo implantation (*61*). The expressions of *Lif* and *Lifr* were significantly reduced in that mouse model, which is not in agreement with our *Zip10* cKO findings. Our present results support their conclusion more precisely: a constitutive expression of epithelial PGR inhibits the detachment of the luminal epithelium. The absence of *Hand2* in uterine tissues was described in a study of embryo implantation failure due to the persistent proliferation of uterine epithelial cells during early pregnancy (*44*). In *Zip10* cKO mice, epithelial cell proliferation had ceased at the timing of embryo implantation (Fig. 3B), suggesting that (at least) the stromal P4-PGR signal that regulates epithelial cell proliferation, such as the FGF-FGF receptor axis (FGFR) (*56*) is likely unimpaired. In contrast, cell proliferation of the subluminal stromal region was enhanced in *Zip10* cKO mice (Fig. 3B, Suppl. Fig. S4), suggesting that stromal cells progressed in the direction of proliferation rather than differentiation.

Despite the higher expression of *Ihh* in *Zip10* cKO mice, we observed that downstream targets of the stromal P4-PGR axis were substantially decreased. We suggest that an epithelial-stromal signal interaction exists to control the epithelial P4-PGR signal. FOXO1, a member of the forkhead transcription factor family, was reported to regulate the epithelial PGR expression (*48*). Ablation of uterine FOXO1 expression led to the PGR expression being retained in the uterine epithelium during the window of receptivity, which resulted in infertility (*48*). In the present study, the FOXO1 expression in the *Zip10* cKO group did not disappear during embryo implantation, indicating that other factors may be involved in the regulation of PGR expression. It remains possible that a lack of ZIP10 affects the functional loss of FOXO1.

*Epas1*, also called hypoxia inducible factor 2α (*Hif2α*), regulates embryo invasion through basement-membrane remodeling and the degradation of the luminal epithelium, since a uterine- and stromal-deletion of *Hif2α* resulted in implantation failure and pregnancy loss (*49*, *50*). Since *Epas1* expression was not reduced in *Zip10* cKO uteri (Suppl. Fig. S6), the loss of *Zip10* in the stroma is not directly linked to the loss of HIF. Our present findings demonstrated that the localization of intracellular GLI1 was highly affected by the intracellular zinc depletion caused by ZIP10 loss in the cultured uterine stromal cells (Fig. 5D,E). GLI1 is a member of the Kruppel family of zinc finger proteins, which are stabilized by zinc ion (*62*). It would be interesting to investigate the relationship between the function(s) of GLI1 and zinc depletion.

In conclusion, we have shown that uterine ZIP10, which functions as an influx of zinc ions into the intracellular space, is essential for successful embryo implantation and placentation in mice. Zinc signal plays an important role in the P4-PGR signal that helps embryo invasion into the endometrium. Zinc deficiency has been a problem in developing countries, but is also present in developed countries where adequate nutrition is available (*63*). A correlation between serum zinc concentrations and birth weight and pregnancy outcomes has been suggested (*63*, *64*). Our results highlight an essential zinc-related molecular mechanism to support the establishment and maintenance of pregnancy.

## Acknowledgements

We thank members of the Laboratory of Animal Reproduction for technical help. We also thank for RIKEN BRC for providing *Zip10^flox^*mice. We also thank Dr. FC. Yang (University of California) for providing *Pgr^Cre^* mice, and Dr. SK. Dey (Cincinnati Children’s Hospital Medical Center) and Dr. Y. Hirota (Tokyo University) for providing *Ltf^iCre^* mice.

---

**Suppl. Fig. S1.** The experimental procedure for the generation of uterine-specific *Slc39a10/Zip10*-deficient mice. **(a)** The genomic DNA region that was deleted to produce *Zip10* conditional knockout (cKO) mice by a Cre-lox system and identified to genotype. **(b)** The breeding strategy to produce *Zip10* cKO females.

**Suppl. Fig. S2.** The morphology of ovaries from the control and *Zip10* cKO mice revealed by hematoxylin and eosin (H&E) staining. Scale bar: 500 µm. CL: corpus luteum.

**Suppl. Fig. S3.** Exogenous hormone control in ovariectomized pregnant females. **(a)** The artificial delayed implantation mouse model. Ovariectomy (OVX) and P4 supplementation with a silicone implant were performed on D4, before the normal embryo implantation process. A single E2 (25 ng/head) injection on D6 induces embryo implantation in this model. Females are euthanized and dissected on D9. **(b)** Representative images of the gross morphology of the uterus from control and *Zip10* cKO mice. *Arrowheads:* the implantation sites. Scale bar: 1 cm. **(c)** Representative images of uterine cross-sections stained by H&E. Scale bar: 500 µm. em: embryo.

**Suppl. Fig. S4.** Subluminal stromal cell proliferation ratio between the control and *Zip10* cKO uteri on D4.

**Suppl. Fig. S5.** COUP-TFII expression in the pregnant uterus. Immunofluorescence of COUP-TFII in sections of implantation sites on D5 from control and *Zip10* cKO mice. Scale bar: 100 µm. st: stroma, em: embryo.

**Suppl. Fig. S6.** *Epas1* mRNA expression of the control and *Zip10* cKO uteri on D6.

