## Supplemental Materials for "Endometrial zinc transporter *Slc39a10/Zip10* is indispensable for progesterone responsiveness and successful pregnancy in mice"

Fig. S1.

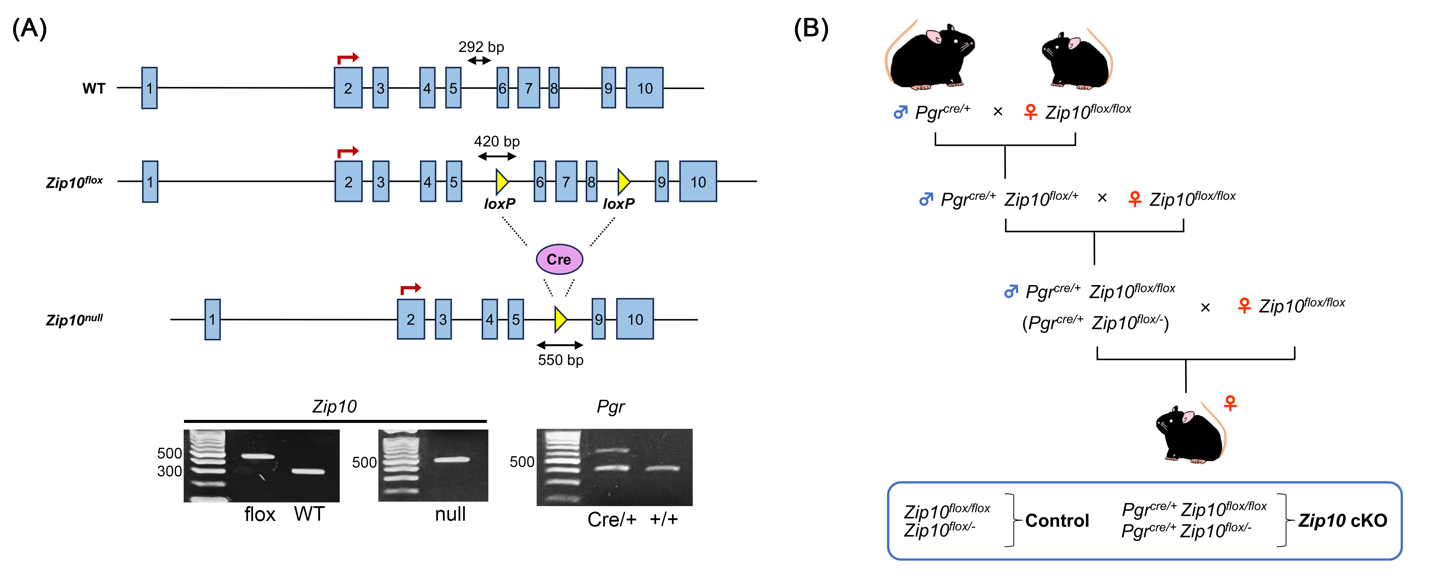

**The experimental procedure for the generation of uterine-specific *Slc39a10/Zip10*-deficient mice.** **(A)** The genomic DNA region that was deleted to produce *Zip10* conditional knockout (cKO) mice by a Cre-lox system and identified to genotype. **(B)** The breeding strategy to produce *Zip10* cKO females.

Fig. S2.

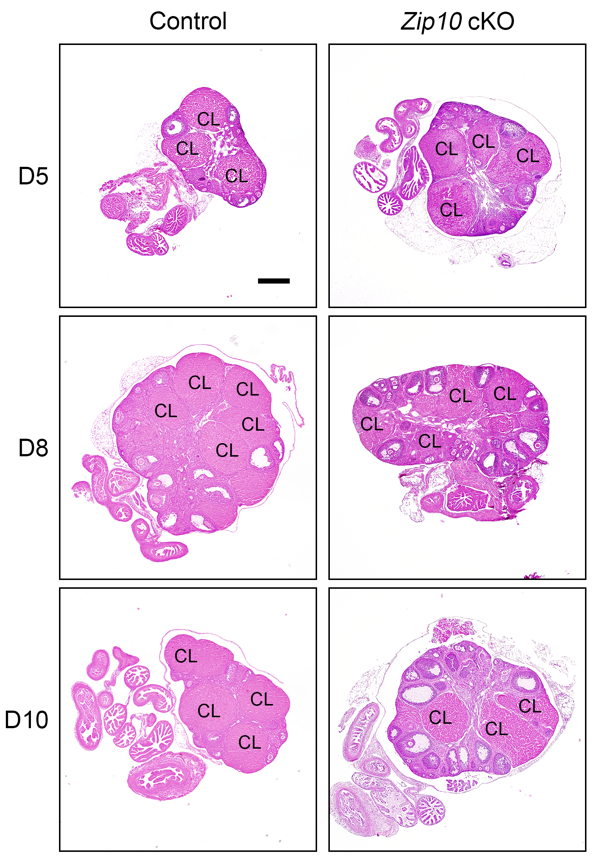

**The morphology of ovaries from the control and *Zip10* cKO mice revealed by hematoxylin and eosin (H&E) staining.** Scale bar: 500 µm. CL: corpus luteum.

Fig. S3.

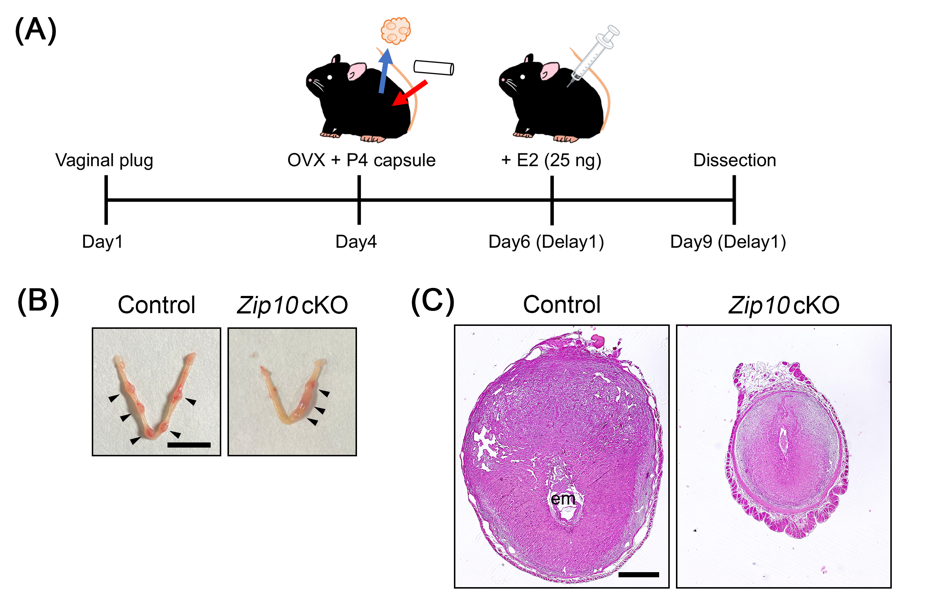

**Exogenous hormone control in ovariectomized pregnant females.** **(A)** The artificial delayed implantation mouse model. Ovariectomy (OVX) and P4 supplementation with a silicone implant were performed on D4, before the normal embryo implantation process. A single E2 (25 ng/head) injection on D6 induces embryo implantation in this model. Females are euthanized and dissected on D9. **(B)** Representative images of the gross morphology of the uterus from control and *Zip10* cKO mice. *Arrowheads:* the implantation sites. Scale bar: 1 cm. **(C)** Representative images of uterine cross-sections stained by H&E. Scale bar: 500 µm. em: embryo.

Fig. S4.

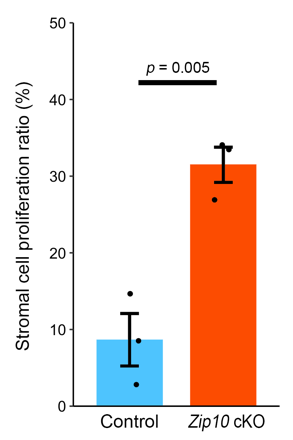

**Subluminal stromal cell proliferation ratio between the control and *Zip10* cKO uteri on D4.**

**Fig. S5.**

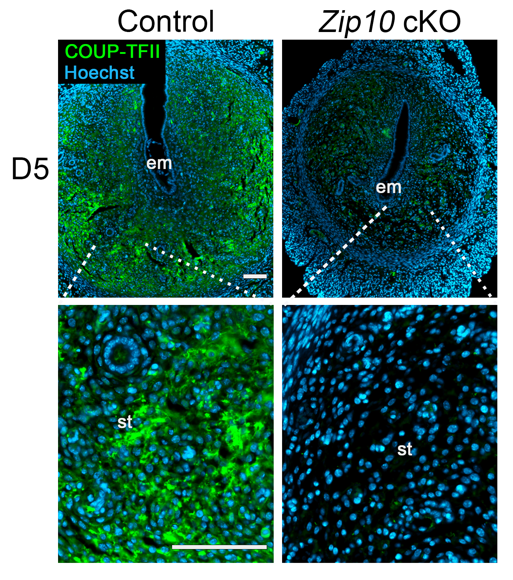

**COUP-TFII expression in the pregnant uterus.** Immunofluorescence of COUP-TFII in sections of implantation sites on D5 from control and *Zip10* cKO mice. Scale bar: 100 µm. st: stroma, em: embryo.

**Fig. S6.**

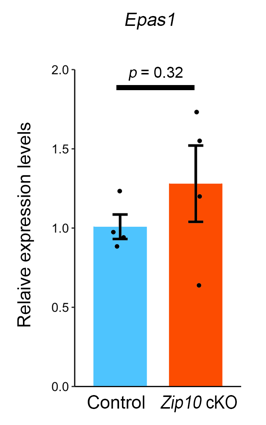

***Epas1* mRNA expression of the control and *Zip10* cKO uteri on D6.**

Table S1. Primary antibody list

| Target | Species | Source | Catalog No. | RRID | Dilution | Application |
| --- | --- | --- | --- | --- | --- | --- |
| ERα | Rabbit | Abcam | ab32063 | AB_732249 | 1:200 | IHC |
| PGR | Rabbit | Abcam | ab101688 | AB_10715248 | 1:200 | IHC |
| MKI67 | Rabbit | Abcam | ab16667 | AB_302459 | 1:200 | IHC |
| FOXO1 | Mouse | Santa Cruz Biotechnology | sc-374427 | AB_10987717 | 1:200 | IHC |
| GLI1 | Mouse | Santa Cruz Biotechnology | sc-515781 | AB_2935677 | 1:200 | IHC/FICC |
| SMO | Mouse | Santa Cruz Biotechnology | sc-166685 | AB_2239686 | 1:200 | IHC |
| CDH1 | Rabbit | Cell Signaling Technology | #3195 | AB_2291471 | 1:200 | IF |
| COUP-TFⅡ | Mouse | Perseus Proteomics | PP-H7147-00 | AB_2314222 | 1:200 | IHC/IF |
| ZIP10 | Rabbit | Miyai *et al*. (2014) #2 | None | None | 1:200 | FICC |

Table S2. Primer list

| Primer | | 5’ - 3’ |
| --- | --- | --- |
| *Bmp2* | Forward | TGGAAAAGGACATCCGCTCC |
|  | Reverse | TGCCACGATCCAGTCATTCC |
| *Epas1*  *(Hif2α)* | Forward | CTGAGGAAGGAGAAATCCCGT |
|  | Reverse | TGTGTCCGAAGGAAGCTGATG |
| *Esr1* | Forward | CCCGCCTTCTACAGGTCTAAT |
|  | Reverse | CTTTCTCGTTACTGCTGGACAG |
| *Gli1* | Forward | GCTTCGGCCAATCACAAATC |
|  | Reverse | CCTGTTTACTCCCACGGTGAA |
| *Hand2* | Forward | GAGAACCCCTACTTCCACGG |
|  | Reverse | GACAGGGCCATACTGTAGTCG |
| *Hoxa10* | Forward | GCCCCTTCAGAAAACAGTAAAG |
|  | Reverse | AGGTGGACGCTACGGCTGATCTCTA |
| *Ihh* | Forward | CTCAGACCGTGACCGAAATAAG |
|  | Reverse | CCTTGGACTCGTAATACACCCAG |
| *Lif* | Forward | GCTGAGCTCTATGAACCAGATC |
|  | Reverse | TGTTCTAGACGCTAGAAGGCC |
| *Lifr* | Forward | GCCTCATTTCTCCGGTTACA |
|  | Reverse | CGAGCACCACTTTGTCTTGA |
| *Nr2f2* | Forward | TCAACTGCCACTCGTACCTG |
|  | Reverse | CCATGATGTTGTTAGGCTGCAT |
| *Pgr* | Forward | CTCCGGGACCGAACAGAGT |
|  | Reverse | ACAACAACCCTTTGGTAGCAG |
| *Ptch1* | Forward | AAAGAACTGCGGCAAGTTTTTG |
|  | Reverse | CTTCTCCTATCTTCTGACGGGT |
| *Wnt4* | Forward | ACACGTGCAGAAACTCAAAG |
|  | Reverse | GAAACTGGTATTGGCACTCCT |
| *Gapdh* | Forward | AGGTCGGTGTGAACGGATTTG |
|  | Reverse | TGTAGACCATGTAGTTGAGGTCA |
